# Unravelling arthropod movement in natural landscapes: small-scale effects of body size and weather conditions

**DOI:** 10.1101/2023.09.25.559245

**Authors:** Garben Logghe, Charlotte Taelman, Florian Van Hecke, Femke Batsleer, Dirk Maes, Dries Bonte

## Abstract

1. Arthropod movement has been noticeably understudied compared to vertebrates. A crucial knowledge gap pertains to the factors influencing arthropod movement at habitat boundaries, which has direct implications for population dynamics and gene flow. While larger arthropod species generally achieve greater dispersal distances and large-scale movements are affected by weather conditions, the applicability of these relationships at a local scale remains uncertain. Existing studies on this subject are not only scarce but often limited to a few species or laboratory conditions.
2. To address this knowledge gap, we conducted a field study in two nature reserves in Belgium, focusing on both flying and cursorial (non-flying) arthropods. Over 200 different arthropod species were captured and released within a circular setup placed in a resource-poor environment, allowing quantification of movement speed and direction. By analysing the relationship between these movement variables and morphological (body size) as well as environmental factors (temperature and wind), we aimed to gain insights into the mechanisms driving arthropod movement at natural habitat boundaries.
3. For flying species, movement speed was positively correlated with both body size and tailwind speed. In contrast, movement speed of cursorial individuals was solely positively related with temperature. Notably, movement direction was biased towards the vegetated areas where the arthropods were originally caught, suggesting an internal drive to move towards suitable habitat. This tendency was particularly strong in larger flying individuals and under tailwind conditions. Furthermore, both flying and cursorial taxa were hindered from moving towards the habitat by strong upwind.
4. In conclusion, movement speed and direction at patch boundaries are dependent on body size and prevailing weather conditions, and reflect an active decision-making process.

## 1. Background

Movement is a crucial component of the life history of many organisms. Across small and large spatial scales, it mitigates the finding of resources and mates, ultimately shaping species distributions and interactions (Bowler & Benton, 2005; Gilbert et al., 1998; McIntyre & Wiens, 1999). Studies involving animal movement tend to focus on vertebrates, leaving other groups such as arthropods severely underrepresented, despite the latter constituting a substantial portion of the global biodiversity (Clark & May, 2002; Kissling et al., 2014). Admittedly, studying arthropod movement involves many inherent challenges, such as their small size and vast species diversity. Nevertheless, understanding the ecological dynamics of arthropods is crucial, as they play pivotal roles in ecosystem functions such as pollination, nutrient cycling and trophic interactions (Yang & Gratton, 2014).

Natural systems are often composed of mosaic landscapes, in which the area is heterogeneously divided into habitats that widely differ in type and suitability (Forman, 2014). Within such landscapes, arthropods may encounter challenging environments while seeking resources or evading unfavourable conditions (Lima & Zollner, 1996; Zollner & Lima, 1999). These unfavourable zones are known as “matrix”, whereas suitable areas are considered “habitat” (Ricketts, 2001; Shreeve & Dennis, 2011). Movement and orientation behaviours at habitat-matrix boundaries will influence the fate of each individual: drifting further in the matrix and losing any connection with suitable environment, such as the home patch or any other potential habitat, or returning to the home patch and remaining philopatric. Unlike routine movements within a habitat, compensatory movement (Peterman et al., 2014; Schtickzelle et al., 2006) through the matrix tends to be rapid with a higher degree of directionality (Hein et al., 2003; Mola et al., 2020; Turlure et al., 2011; Van Dyck & Baguette, 2005). Consequently, it can be expected that crossing a hostile matrix will impose substantial dispersal costs (Bonte et al., 2012), particularly for small-bodied organisms like arthropods (Hillaert et al., 2018). Behaviours at boundaries are therefore expected to have pervasive effects on population dynamics, gene flow and hence the adaptive capacities of species in fragmented landscapes (Batsleer et al., 2023; Driscoll et al., 2013).

Body size stands out as one of the most relevant factors influencing arthropod movement and their capacity to orientate, due to its association with various morphological, physiological and life-history traits (White et al., 2007; Woodward et al., 2005). Numerous arthropod taxa, ranging from cursorial groups (i.e. individuals running on the ground) like ants and beetles, to flying taxa such as butterflies, bugs and dragonflies, seem to consequently exhibit an allometric relationship between body size and mobility (Hirt et al., 2017; Hurlbert et al., 2008; Kuussaari et al., 2014; Sekar, 2012; Wootton, 2020). Larger arthropod individuals, both within and across species, generally display greater mobility, resulting in higher movement speeds and larger travelling distances.

Besides species-specific characteristics such as body size, arthropod movement can also be influenced by external conditions, such as weather. The impact of local weather conditions should be most apparent for small-scale movement bouts, because of the high variability of weather conditions at small temporal and spatial scales. A first component of local weather conditions is the prevailing temperature. Higher ambient temperatures have been shown to positively impact the activity and running speed of invertebrate ectotherms, owing to improved oxygenation rates and enhanced kinetic muscle properties (Frizzi, 2018; Hurlbert et al., 2008). Moreover, arthropod species often adjust their movement behaviour according to the local thermal conditions, with some even requiring a minimum ambient temperature to initiate movement (Frizzi, 2018; Knight et al., 2019; Knoblauch et al., 2021). However, movement responses to increased temperatures are also constrained by critical thermal upper limits, leading to impaired movement capacity when temperatures exceed these thresholds (Chown & Nicolson, 2004; Dudley, 2000; Heinrich, 1995). A second component of local weather is wind. The direction and force of the wind exert considerable influence on movement distances and decisions, especially for flying insects undertaking long-distance dispersal (Knoblauch et al., 2021). Additionally, many arthropods possess anemotactic reception, allowing them to accurately estimate the direction of prevailing winds by sensing air currents, and consequently, make informed choices regarding movement trajectories (Buehlmann et al., 2020; Knoblauch et al., 2021).

Most of the previously mentioned research on the relationships between arthropod movement, body size and weather conditions focuses on large-scale distances, and usually involves only a limited number of large and highly mobile species (Frick et al., 2013; Knight et al., 2019; Knoblauch et al., 2021). Small-scale movement studies investigating biometeorological relationships in arthropod movement are scarce (Aikman & Hewitt, 1972; Eguagie, 1974; Merckx et al., 2008) and especially lacking when considering behaviour at habitat boundaries. Traditional methods, such as capture-mark-recapture or tracking devices, have limitations in achieving large sample sizes, often leading to a narrow focus on individual species and hindering generalizations across taxa and life history traits (Goodwin & Fahrig, 2002; Jonsen et al., 2001). Moreover, concerns have been raised about the use of such tracking devices, as the impact of these tags on arthropods is often neglected, ultimately biasing data (Batsleer et al., 2020). While some notable efforts have been made to study arthropod exploratory movement through image-based tracking under laboratory conditions, the applicability of these findings to natural field conditions remains uncertain (Hirt et al., 2017; Terlau et al., 2023).

This study aims to understand the interplay between movement, body size and weather conditions at habitat boundaries by conducting a field release experiment on a diverse range of arthropod species. In contrast to macro-ecological approaches that treat dispersal as colonisation at biogeographical scales (Weil et al., 2023), our methodology focuses on individual departure and settlement behaviour. Multi-species experiments under similar environmental settings have the potential to yield generalisations on trait-environment correlations across and within species, and to understand potential evolutionary determinants of these relationships (Felsenstein, 1985).

Two different nature reserves consisting of dune and heathland habitats were used as field sites, which differed in spatial heterogeneity. This allowed us to assess whether arthropods choose to move towards suitable habitat areas when venturing in the matrix. The realised movement speed and direction of arthropods at the patch boundary—without relying on tracking devices—were measured and correlated with individual body size and prevailing weather conditions. Movement behaviours after release represent movements that we consider to be relevant for short-term displacements when organisms leave their habitat and enter the matrix. We expect (i) a positive correlation between movement and body size, (ii) the strengthening of this effect under warmer temperatures and stronger tailwinds, (iii) an influence of wind conditions on the movement direction of flying insects, and (iv) a biased movement direction towards suitable environments due to the perception (correct interpretation by the individual) of the environment and avoidance of risks when crossing the matrix.

## 2. Methods

Permits to access the field sites and capture protected species were granted by the Agency of Nature and Forest. This study did not require ethical approval.

### 2.1 Experimental fieldwork

The fieldwork for this study was conducted in the summer of 2021 at two locations in Belgium. The first site was a coastal dune grassland in De Panne, while the second site consisted of heathland patches in Kalmthout (Appendix S.1). Both sites differed in structural heterogeneity of the landscape. In the coastal dune site at De Panne, large areas of open sand (representing the matrix) bordered large areas of shrubland, thus creating a “binary” landscape. In Kalmthout, patches of open sand were considerably smaller, and were embedded within a dense heath landscape, thereby creating a patchier landscape where matrix and habitat surrounded each other. By selecting these two distinct sites, we aimed to determine whether arthropod movement direction at habitat boundaries is either random or biased towards vegetated areas. If arthropods indeed show an internal drive to move towards habitat patches, we should find a strong bias towards one direction in the “binary” landscape at De Panne, while movement should be more random in the patchy landscape at Kalmthout (Appendix S.1).

Within the vegetated areas at both sites, arthropod individuals from various species were randomly captured either by hand or by sweep netting. Each captured individual was assigned a unique ID and photographed in the field with a graph paper (1 mm squares) as background. To assess the movement behaviour of these arthropods, separate set-ups were designed for flying and cursorial (i.e. running on the ground) individuals, as we anticipated different movement speeds and perceptual abilities. In a large area of open sand, which functions as matrix for species living in vegetated areas, two circular set-ups were demarcated by placing coloured flags at five different radii with uniform interval distances (2 metres for flying individuals and 20 centimetres for cursorial individuals). The shortest distance from the point of release to habitat measured 1 m and 10 m for respectively the experiments on cursorial and flying movement (west orientation for De Panne and northwest orientation for Kalmthout). In accordance with the earlier described habitat heterogeneity, distance to habitat was more than 100 m in other cardinal directions for De Panne, while for Kalmthout these distances were less than 50 m.

Movement speed was measured using a chronometer, whereby the movement time was recorded as the duration an individual took to traverse either 10 meters (for flying locomotion) or 1 meter (for cursorial locomotion), or when an individual came to stop at a specific distance. The directions of movement and wind were categorised using cardinal directions (N, NE, E, SE, S, SW, W or NW), and then converted into angles in degrees, with west serving as the zero degrees reference (see further). By comparing the movement and wind angles, we were able to determine whether the movement was aligned with the wind or not, and to differentiate between movement upwind and movement assisted by tailwinds. Specifically, we defined tailwind movements when the direction of movement deviated by a maximum of 45 degrees from both sides of the prevailing wind direction, while upwind movements deviated by at least 135 degrees from both sides of the prevailing wind direction (or 45 degrees from both sides of the direction of the tail; Figure 1). West was chosen as the zero-degree reference, as the vegetated areas at De Panne were located in this direction relative to the experimental set-up (Figure 1). This decision assumed that the arthropods would exhibit a preference for returning to the location where they were initially caught, since they were released in a matrix with very few available resources. The surface area of the matrix was sufficiently large to assume that moving through it would bring a high energetic cost and uncertainty for the released arthropod. By adopting the orientation as shown in Figure 1, we aimed to gain insights in navigation behaviour, as movement patterns would likely reflect the tendency and ability to return to the habitat of origin.

**Figure 1:**
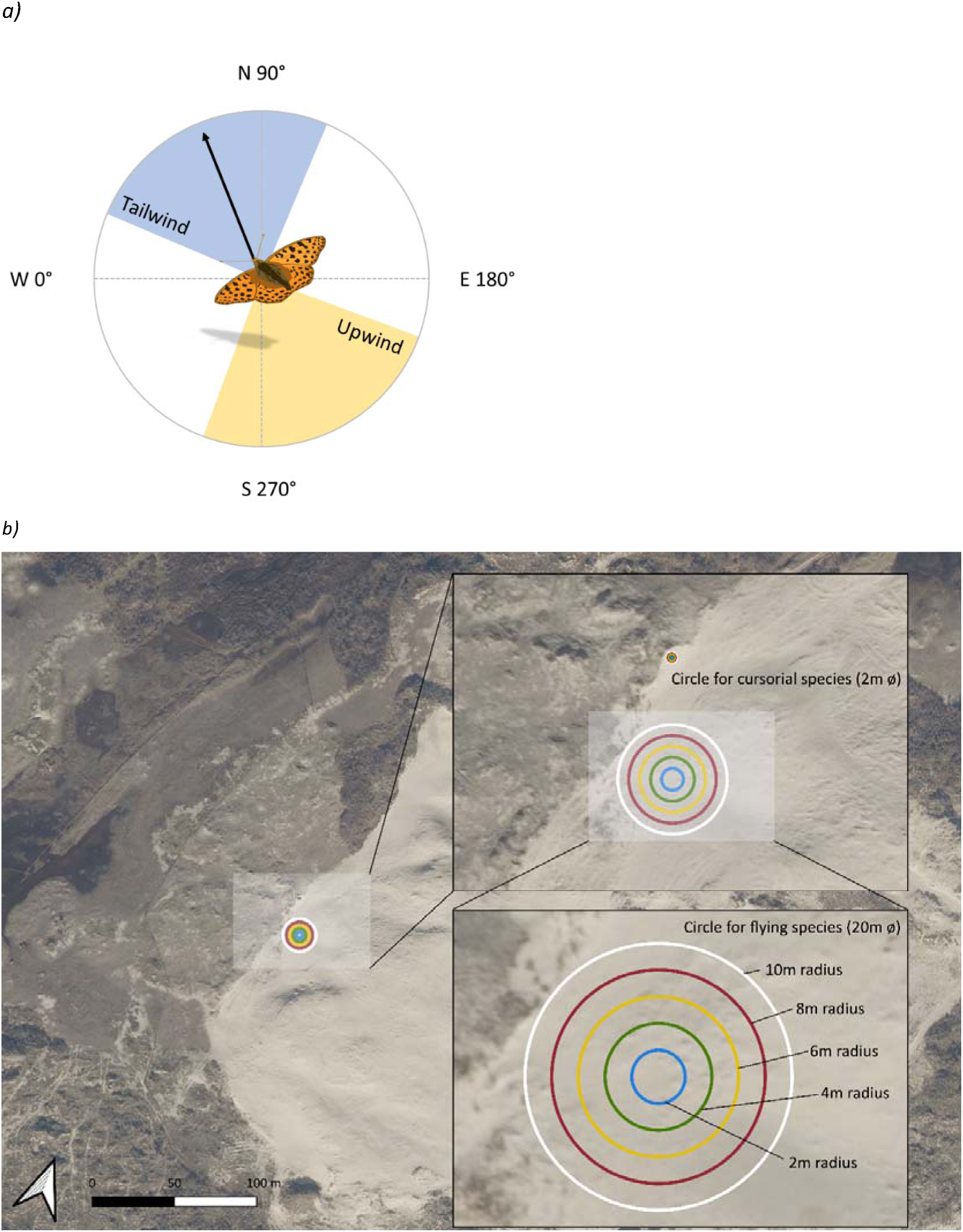
Schematic representation of how the cardinal directions were converted into angles. The black arrow indicates the direction of the movement. The blue area indicates all wind directions that can be classified as tailwind in this specific example, while the yellow area indicates upwind. b) Map of the experimental set-up at De Panne, with the vegetated areas to the west. Circular setups were drawn at a realistic scale. The illustrations in this figure were created by author Charlotte Taelman, aerial photograph (winter 2022) obtained from Agency for Information Flanders (geopunt.be).

During measurements, the captured arthropods were temporarily housed in dark boxes to minimize stress and prevent mortality due to overheating. Each individual spent a maximum of thirty minutes in these boxes. Subsequently, the arthropods were taken out one by one and released in the open sand. To mitigate the likelihood of an escaping movement response and directional bias arising from the researchers’ presence, cursorial animals were released at a distance of 1 meter from the observer, whereas flying animals were released above the head of the observer. Individuals were not chased but followed by eye, and timed from the moment of release until the moment they reached the edge of the experimental arena. Every individual had complete freedom of where to move and when to stop. By doing this, we aimed to minimize stress and present the option to actively choose movement directions.

Every half-hour of the release experiments, wind direction, wind speed and temperature were recorded at an average height of 1.5 to 1.8 metres, using a weather station equipped with a built-in thermometer and anemometer. Wind speed was calculated as the average of the minimum and maximum values measured within a half-minute interval. Sampling days were carefully selected to ensure suitable weather conditions for representative arthropod movement: a cloud cover below 60% and temperatures between 15°C and 34°C.

Due to practical constraints, determining the dry weight of individual arthropods was not feasible. Instead, body length was used as an alternative proxy for body size (Ganihar, 1997; Martin et al., 2014; Moretti et al., 2017). The pictures of the arthropods taken in the field were analysed using ImageJ version 1.52a (Schneider et al., 2012). The grid pattern in the background of the photos was used as reference to measure the length of each individual’s body, starting from the cephalon (excluding the antennae) and extending to the tip of the abdomen.

### 2.2 Statistical analyses

All data analyses were conducted using R version 4.2.2 (R Core Team, 2021). The “dplyr” package (Wickham et al., 2022) was used for general data manipulation and organization. All visuals were created with the “ggplot2” (Wickham, 2016), “RcolorBrewer” (Neuwirth, 2022) and “showtext” (Qiu, 2023) packages.

#### 2.2.1 Movement speed

As a first explorative analysis, a Principal Component Analysis (PCA) was conducted, separately for flying and cursorial species to understand how species movement behaviours group according to their traits, phylogeny and the prevalent environmental conditions.

To confirm any observed relationships between movement speed, body size and weather conditions in flying and cursorial arthropods, a distinct Bayesian Generalized Linear model was employed for each group (“brms” package (Bürkner, 2021)). The dependent variable in these models was movement speed, while the independent variables were body size, temperature and wind speed (all standardized to a mean of zero and a standard deviation of one). The models were fitted with a Gaussian distribution using 4 chains and 4000 iterations to ensure reliable results. Non-informative flat priors were used for the model parameters. As we expected wind speed to have opposing effects depending on its orientation relative to an individual’s movement, we constructed additional models by splitting the dataset into individuals that moved either tail- or upwind and ran two separate models. This approach was used to test potential differences in responses and proved to be more efficient than the incorporation of an interaction between two circular variables to the Bayesian models.

Since the models did not account for the potential influence of phylogenetic relationships between species, additional models were constructed to address this issue. For this purpose, phylogenetic trees were created by linking the species names to the Open Tree Taxonomy database, establishing a tree based on the known phylogenetic relationships between species (“rotl” package (Michonneau et al., 2016)). Branch lengths were calculated with the “ape” package (Paradis & Schliep, 2019) using the “Grafen” method (Grafen, 1989), leading to the construction of a phylogenetic covariance matrix (“geiger” package (Matthew et al., 2014)). First of all, this tree was used to calculate Pagel’s lambda, a measure for the phylogenetic signal in a specific trait, for movement speed and direction of both flying and cursorial species (“phytools” package (Revell, 2012)). Secondly, the phylogenetic distances between species were incorporated into all Bayesian Generalized Linear models (similar to those above) as blocking factors. By doing so, the models considered the phylogenetic relationships between species when analysing the correlation between movement speed and the other variables. We contrast insights from the non-corrected models (ecological responses of the sampled arthropod community) and phylogenetically corrected models (ecological responses of arthropod species, corrected for the shared, and hence non-independent evolutionary history of life histories and morphologies). As such, we can understand whether the community-wide movement responses are a direct outcome from the specific reactions of the sampled species, or whether they have evolved as a general pattern across species.

#### 2.2.2 Movement direction

The analysis of arthropod movement behaviour in different wind scenarios involved several circular statistics tests (“circular” package (Agostinelli & Lund, 2022)). These analyses were performed separately for both sampling sites to analyse the effect of structural heterogeneity within the landscapes.

To assess the correlation between wind direction and movement direction, the Jammalamadaka-Sarma correlation coefficient (Jammalamadaka & Sarma, 1988) was calculated for both flying and cursorial arthropods within different taxonomic orders, limited to orders represented by 8 or more samples. Additionally, the Rayleigh’s test of Uniformity (Pewsey et al., 2013) was performed to examine whether there was a unimodal directional bias in the distribution of arthropod movement directions. The significance of the results from these tests was evaluated by correcting for multiple testing using the Benjamini-Hochberg method (Benjamini & Hochberg, 1995), addressing potential issues of false positives.

Furthermore, several Bayesian Generalized Linear Models were fitted to explore the relationship between body size and movement direction. For these analyses, only data from De Panne was used since the difference in direction between habitat and matrix was much clearer compared to the site at Kalmthout. A binary value (0 or 1) was assigned to indicate whether or not an individual had moved towards the habitat from where they were initially caught. Deviations greater than 45 degrees from the zero-degree point (west) were considered as not moving towards the habitat and were assigned a value of 0. Body size and wind speed were used as independent variables, while this binary value was the dependent variable, modelled using a binomial distribution with a logit link function. Temperature was omitted from these models because we considered it irrelevant in the context of navigation. As with movement speed, the datasets were again split to test whether different responses were present under upwind or tailwind conditions. As for movement speed, each model was also corrected for phylogenetic relationships, which involved constructing phylogenetic trees using Open Tree Taxonomy and incorporating phylogenetic distances as blocking factors in the models.

## 3. Results

In total, 1355 arthropod individuals from 255 different species were tested (Appendix S.2). Among these individuals, 762 exhibited flying locomotion, while the remaining 593 displayed cursorial locomotion. Measured temperatures ranged from 15°C to 40°C, while wind speed varied between 0 and 6.4 metres per second.

### 3.1 Movement speed

The PCA shows Odonata, having larger body sizes, to be separated from the other studied arthropod orders (Figure 2). For both cursorial and flying arthropod groups, PC1 was correlated with temperature and windspeed and PC2 with body size (Appendix S.3). That the first axis explains respectively 36% of the variation in flying species and 34% of the variation in cursorial species, shows that weather conditions are indeed an important determinant of movement. We further explore interspecific variation in more detail by means of (phylogenetically corrected) linear models.

**Figure 2:**
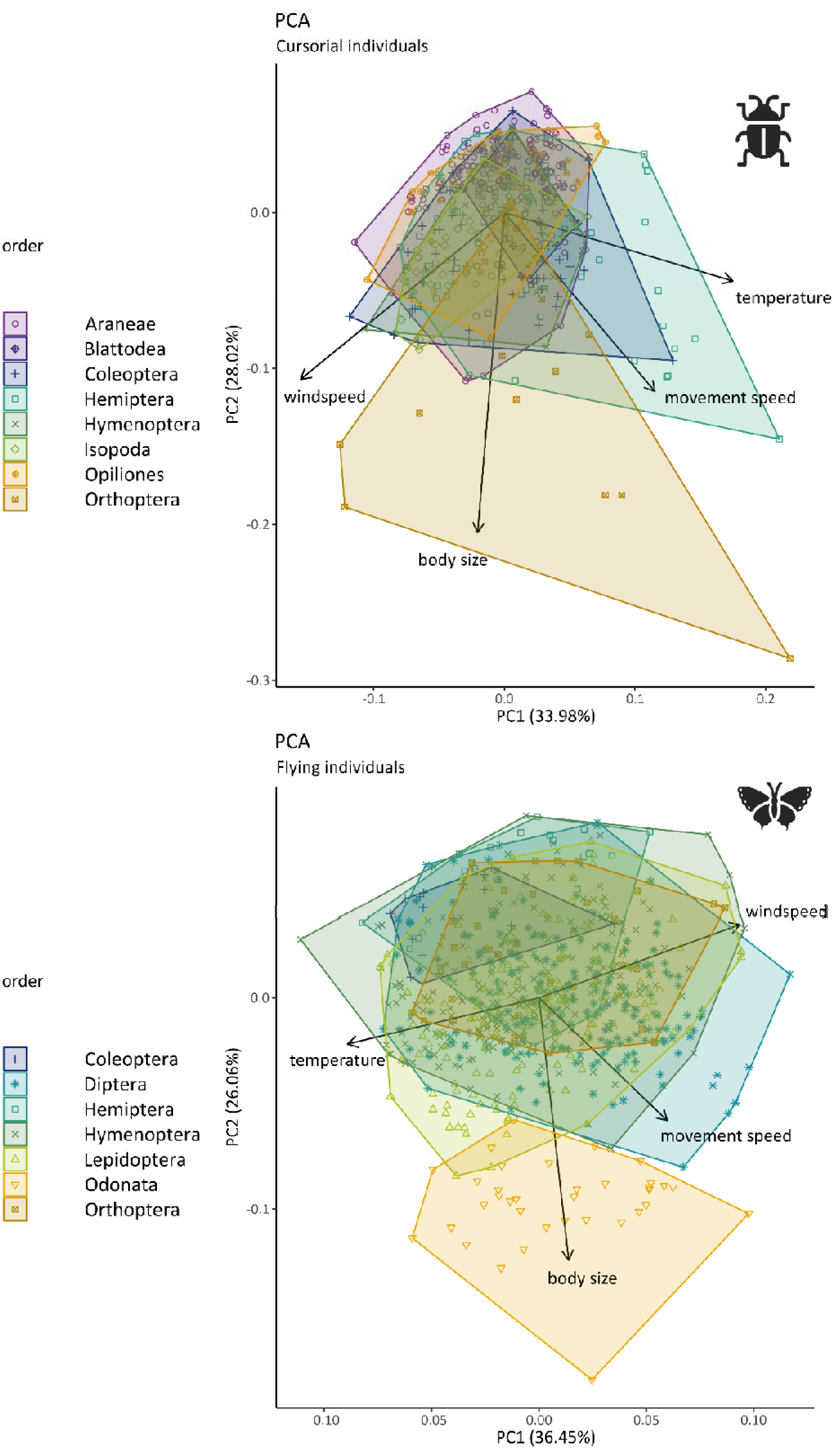
PCA results for flying (top) and cursorial (bottom) individuals. Dots represent individuals with symbol colour and shape according to their taxonomic order. A convex polygon is drawn around individuals of the same taxonomic order, by connecting the most extreme values. The arrows represent the direction and magnitude of each variable’s contribution to the principal components. Longer arrows indicate stronger influence, while the direction shows the correlation between variables and components. The angle between the arrows reflects the correlation between traits; very small angles indicate strong positive correlation, while very large angles indicate strong negative correlation.

In flying species, large species generally exhibited higher movement speeds compared to smaller ones (Figure 3a, left). However, this relationship appears to reach an optimal point around 15 mm body size, beyond which a further increase in body size does not lead to higher movement speeds (Appendix S.4.2). The overall positive impact of body size and wind speed on movement speed of flying species remains when only tailwind movements are considered, but disappears when movement occurs upwind (Figure 3b, c, left). For tailwind speed, the correlation also demonstrated a saturation point, which occurred around 3 m/s wind speed, beyond which movement speed did not increase further (Appendix S.4.1). The overall positive effect of wind speed on movement speed is thus primarily driven by the effects found under tailwind conditions (Figure 3, left). Temperature had no specific effect on the movement speed of the studied flying insects (Figure 3, left). Pagel’s lambda of movement speed was notably high (λ=0.733), indicating a strong phylogenetic signal. Nevertheless, as expected from the overlap among taxa in the PCA, phylogenetic corrections had no substantial qualitative impact on the posterior distributions of the effect sizes and their interpretation (Figure 3). Species evolutionary history is thus an important driver of movement speed, independent of the species’ attained body sizes. Since a phylogenetic correction did not affect any of the patterns, responses to weather are shared by all species present in the study area, independent of their phylogenetic background.

**Figure 3:**
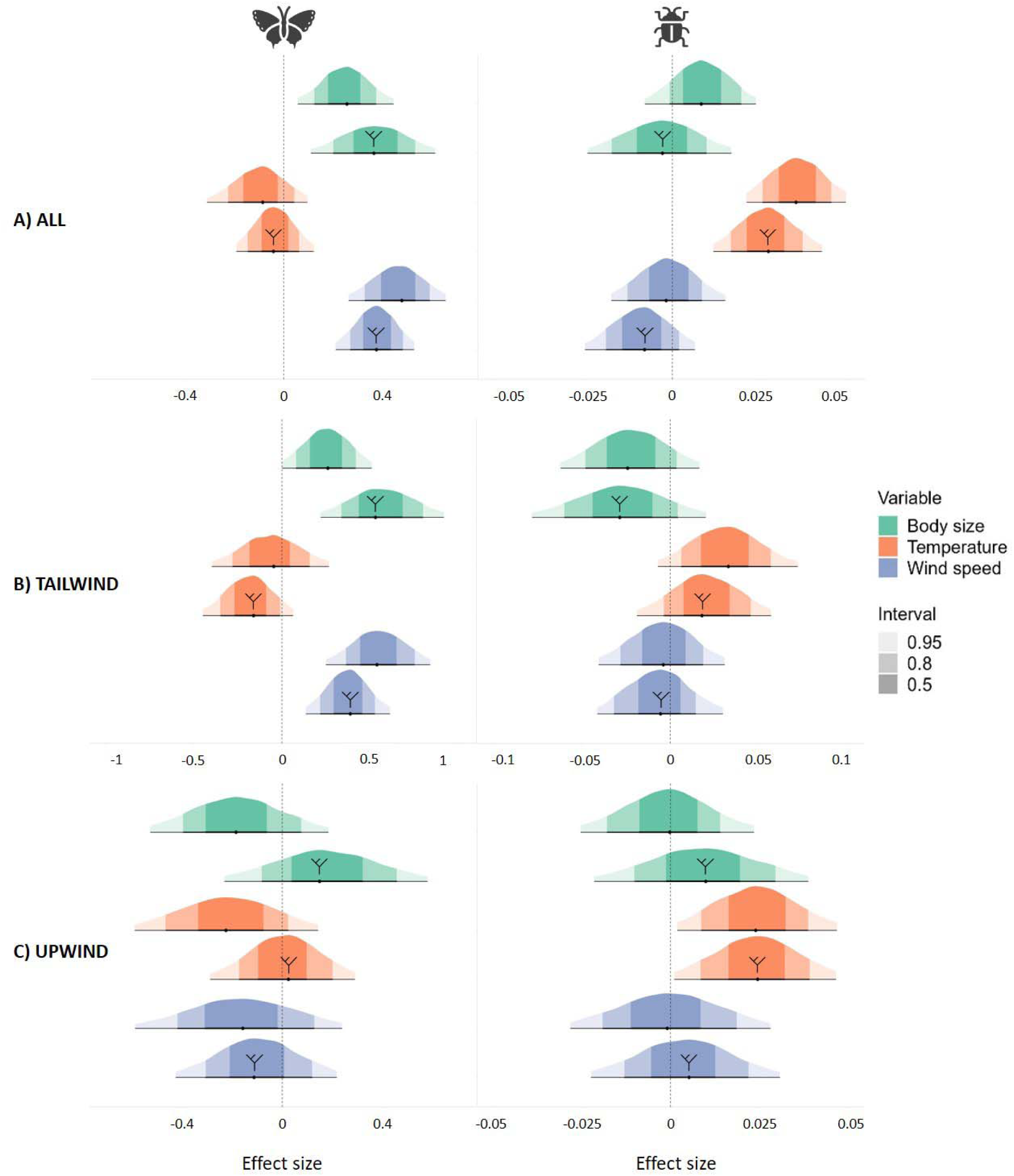
Posterior distributions of the effect sizes between movement speed as dependent variable and body size, temperature and wind speed as independent variables. For a) all wind conditions, b) tailwind conditions (n=527) and c) upwind conditions (n=281). Posterior distributions that are marked with a phylogenetic tree resulted from the phylogenetically corrected models. Effect sizes were deemed clearly determined when 0 was not included in the 95% credibility intervals of the posteriors.

The movement speed of cursorial species was, in contrast, only affected by temperature (Figure 3, right), with the positive effect indicating higher speeds reached under warmer conditions. The Pagel’s lambda of movement speed for the cursorial species was considerably smaller compared to flying species (λ=0.357). Closely related cursorial arthropods are therefore more variable in their movement speed relative to their flying counterparts. For a detailed overview of the results from these models, see Appendix S.5.1.

### 3.2 Movement direction

Arthropod movement exhibited a noticeable directional bias, indicating that individuals did not move randomly (Figure 4). The R-values, representing concentration of movement and thus directional bias, were significant after correction for multiple testing for most taxonomic orders (Appendix S.6). In addition, the Jammalamadaka-Sarma correlation coefficients between wind and movement direction did not show a general pattern across taxa, with some orders following the wind more than others (Appendix S.7). However, these correlations where only significant for Lepidoptera (Kalmthout, *r*=0.36), Araneae (De Panne, *r*=0.34) and Diptera (De Panne, *r*=0.19).

At De Panne, where the habitat of the arthropod species was clearly located in one direction (west), movement direction was mostly oriented towards habitat areas across all observations (Figure 4.1). The lowest R-value at this site (R=0.19; p=0.05) was associated with a southwest wind opposite to the habitat direction, indicating lack of any directionality of movements under these wind conditions. In contrast, the highest R-value and hence strongest directionality prevailed under southeast winds, which largely aligned with the orientation towards the habitat (R=0.71, p<0.01). Furthermore, the Jammalamadaka-Sarma coefficient across all observations was positive (*r*=0.07), indicating a minor effect of wind direction on movement direction. In contrast, at the Kalmthout site, which was surrounded by suitable habitat in multiple directions, movement directional bias was only significant under NE winds (R=0.44; p<0.01). This indicates that arthropod movement was more random compared to the site at De Panne. Additionally, the Jammalamadaka-Sarma coefficient was larger (*r*=0.14) compared to De Panne, suggesting that movement direction was more positively aligned with wind direction (Figure 4.2).

**Figure 4.1:**
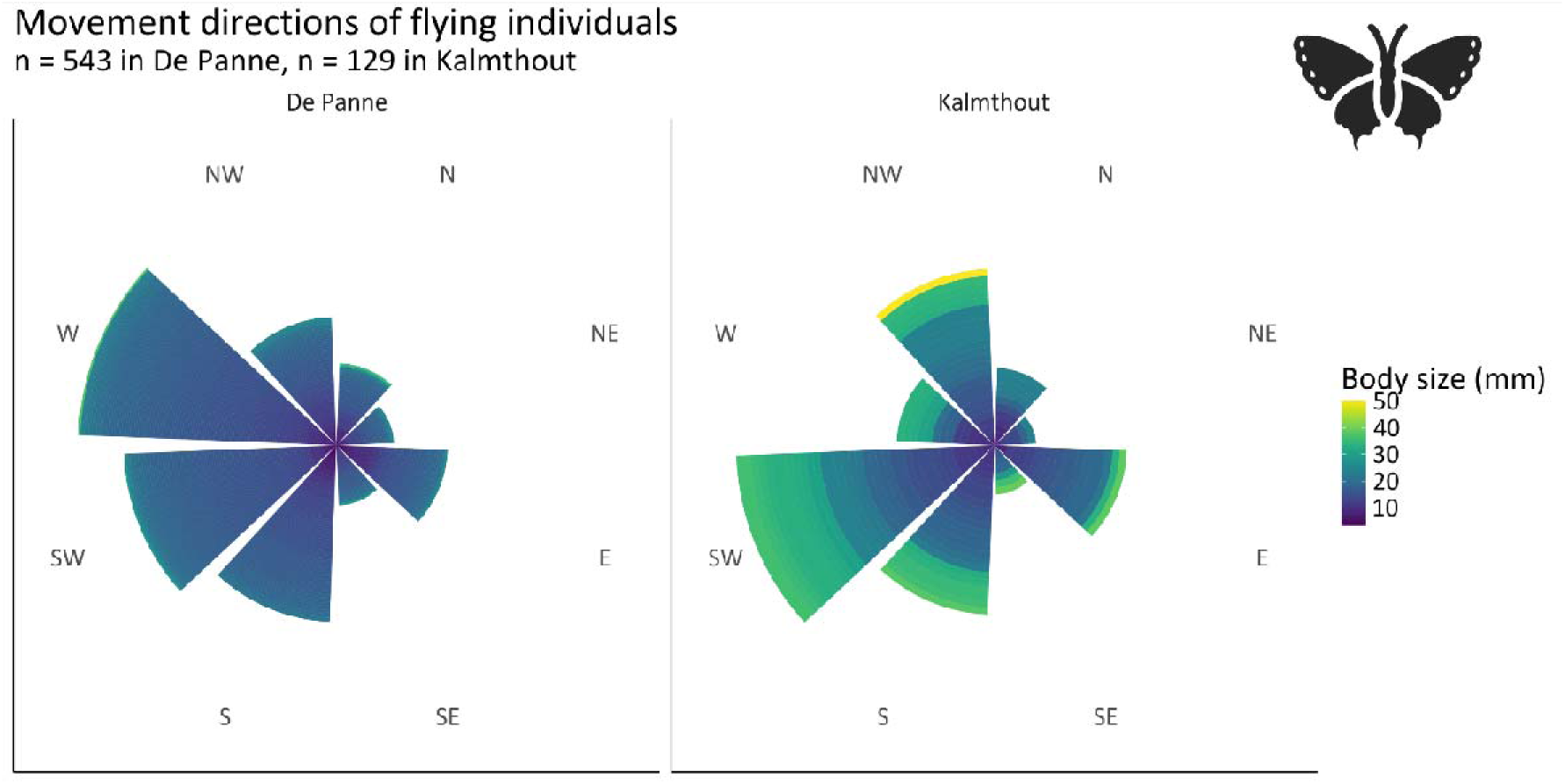
Circular bar plots of movement directions of flying species at both field sites. Each concentric slice on the bar represents an individual and is coloured according to its body size. Thus, larger bars indicate a higher number of individuals that moved in that direction. Similar figures for specific wind directions can be found in Appendix S.8.

**Figure 4.2:**
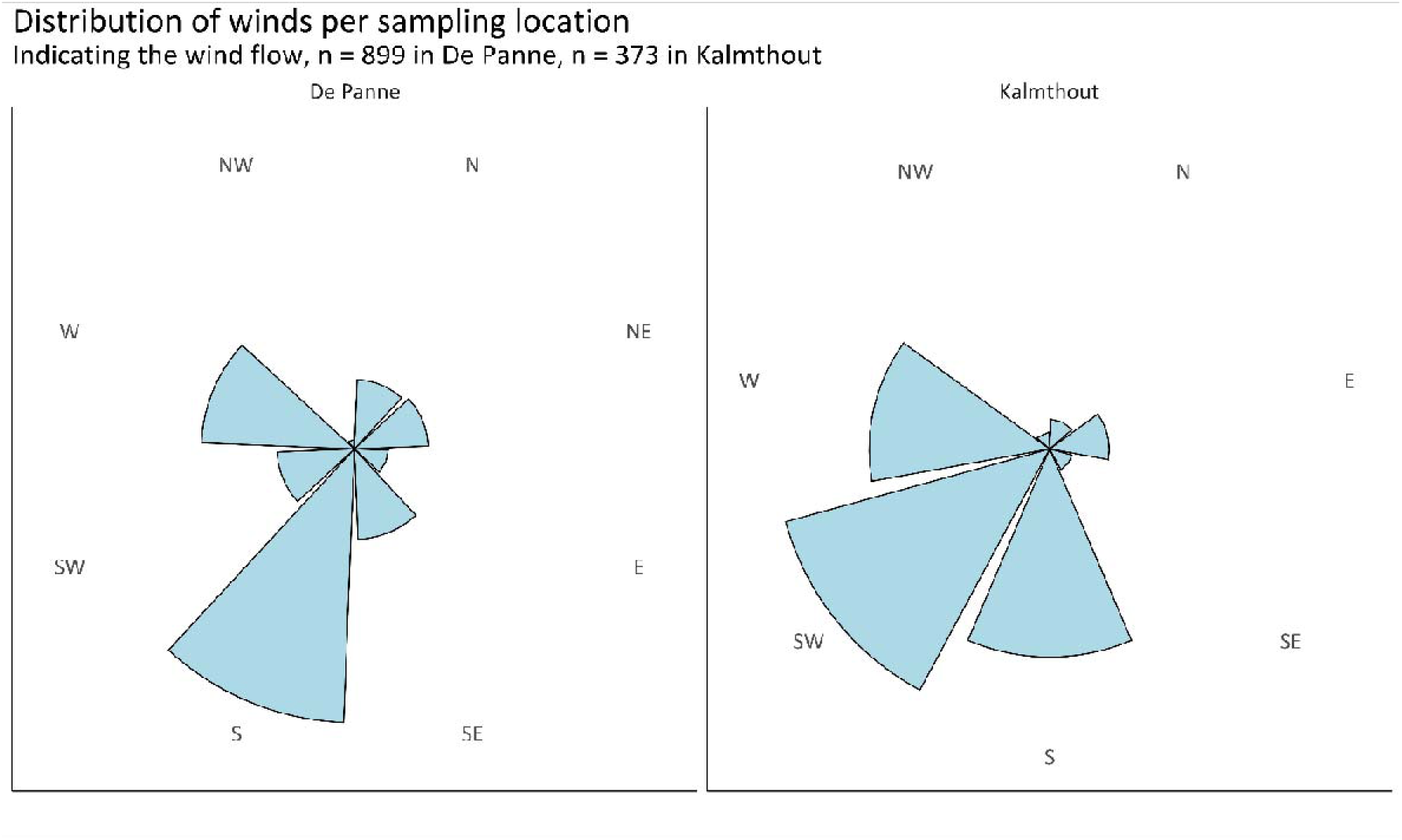
Circular bar plots of dominant wind directions at both field sites. The larger the slices, the more observations were made under that specific wind direction. Wind speed data for each wind direction can be found in Appendix S.9.

Movement direction did not only correlate to wind direction, but additionally to wind speed as well. We ran Bayesian generalized linear models for the experiments in De Panne only, which showed that higher wind speeds reduce the efficiency of flying individuals to move towards habitat (Figure 5a, left). This effect was mainly steered by individuals tested under upwind conditions (Figure 5c, left).

**Figure 5:**
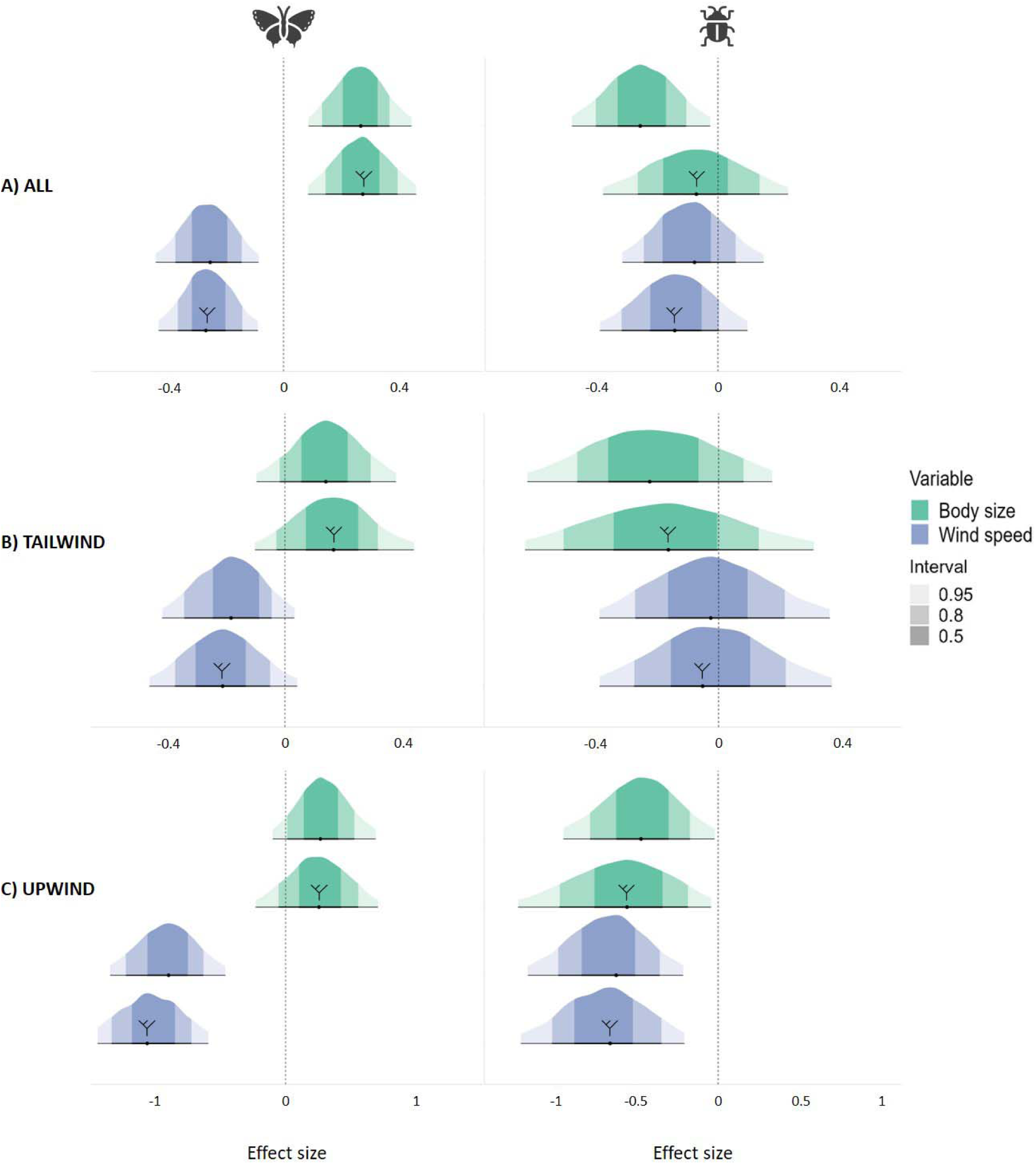
Posterior distributions of the effect sizes between movement direction as dependent variable and body size and wind speed as independent variables. For a) all wind conditions, b) tailwind conditions (n=440) and c) upwind conditions (n=248). Only data from the De Panne site was used for this analysis. Posterior distributions that are marked with a phylogenetic tree resulted from the phylogenetically corrected models. Effect sizes were deemed clearly determined when 0 was not included in the 95% credibility intervals of the posteriors.

Similarly, cursorial species only showed a negative correlation between wind speed and movement direction under upwind conditions (Figure 5, right). This indicates that strong upwinds lower the propensity of both groups to navigate back to the habitat. These patterns remained conserved under phylogenetic correction (λ < 0.001), indicating generally similar arthropod-wide responses. Habitat-directed movements also differed with body size. Larger flying species are more prone to move towards habitat, while the pattern is reversed for cursorial species (Fig 5a, right). Additionally, the effect of wind speed on cursorial navigation only holds under upwind conditions when phylogenetically corrected (λ = 0.210). Our observed community-wide responses were thus biased by the specific sample of tested species. For a detailed overview of the results from these models, see Appendix S.5.2.

## 4. Discussion

We used release experiments to gain insights into the condition-dependency of small-scale arthropod movement at their patch boundary. Movements at this spatiotemporal scale can determine the future trajectories within or back from the matrix in fragmented habitats. These fundamental movement elements are anticipated to eventually shape large-scale canonical and lifetime movement patterns (Getz & Saltz, 2008), and thus patterns of gene flow, colonisation-extinction dynamics and species assembly in fragmented landscapes (Batsleer et al., 2023; Jeltsch et al., 2013). Using a standardised approach under natural conditions, we demonstrate the importance of phylogeny, body size and weather conditions on arthropod movement in the hostile matrix close to the patch boundary. Larger flying arthropods exhibited greater movement speeds compared to their smaller counterparts, while the presence of strong tailwinds further increased their movement speed. In contrast, the movement speed of cursorial arthropods appeared unrelated to body size but increased with temperature. Notably, our observations revealed a clear directional bias towards suitable habitat at both sites, indicating an internal drive among arthropods to move towards suitable environments. However, species showed varying levels of navigational proficiency, with larger flying and smaller cursorial arthropods displaying a greater tendency to move in the direction of suitable habitat. Furthermore, our study identified strong upwinds as hindering the movement of both groups, impeding the capacity for navigation towards suitable habitat.

The positive correlation between movement speed and body size in flying arthropods was in alignment with our expectations. Previous studies already established that—within a taxon—larger arthropods tend to move faster (Kuussaari et al., 2014; Wootton, 2020). Our study emphasizes that this effect remains consistent across multiple taxa, even after accounting for phylogenetic relationships. Consequently, there is a difference in movement speed between taxa, but the positive relationship between movement speed and body size remains conserved. For example, larger bees are generally faster than smaller bees or smaller butterflies, but do not necessarily surpass the pace of a larger dragonfly.

In contrast, the absence of a correlation between movement speed and body size in cursorial arthropods does not match the positive relationship retrieved in previous studies (Foellmer et al., 2011; Hirt et al., 2017; Hurlbert et al., 2008). A first possible explanation is that a relatively limited range of body sizes was tested, spanning from 1.21 mm to 21.31 mm, preventing us from detecting a signal when effects are mainly determined by very large species. Secondly, our proxy for body size could obscure true relationships. For instance, harvestmen may appear small in our dataset due to their compact body length, but their long legs could enable faster running compared to species with similar body length (Arnold et al., 2017; Kaspari & Weiser, 1999; Smith et al., 2012). The use of a relative measure (e.g. leg or wing length relative to body length) remains, however, hard to generalize across taxa and thus limits the applicability for cross-taxa comparisons. A third potential explanation is linked to the interaction between the body mass of an individual and the texture of the surface on which it moves. Sand is a porous substance, in which heavier individuals might sink deeper than smaller ones, hence slowing their movement.

In addition to body size, wind speed also positively influenced the movement speed of flying individuals. Conversely, when individuals faced upwind, the effect reversed. This implies that movements away from habitat boundaries may be more pronounced when arthropods fly with tailwind, which can assist them in crossing the matrix (DiLeo et al., 2022). However, under natural conditions, the prospect of wind blowing arthropods out of their habitat might deter them from departing, as for instance found in ballooning spiders (Bonte et al., 2007). In our study, however, individuals initiated movement within the local matrix after being placed there involuntarily. Release experiments, like those we conducted with many species, or similar work on single species (Bonte et al., 2004; Kallioniemi et al., 2014; Merckx & Van Dyck, 2007; Schultz et al., 2012), represent movement behaviour at habitat boundaries. However, these experiments are less likely representative of routine behaviours such as foraging within suitable habitat, or special dispersal behaviours when crossing the matrix (Bonte et al., 2012; Clobert et al., 2009).

Surprisingly, temperature did not impact the movement speed of flying individuals. This can potentially be attributed to the capacity of many flying arthropods to elevate their body temperature through wing movement (Heinrich, 1974). Moreover, our dataset lacked extreme (hot or cold) temperatures and we used an approximative measure for temperature, measured at the height of the observer, not in the direct surroundings of the individuals. Both can potentially mask any trend between temperature and movement of flying arthropods. In contrast, cursorial individuals demonstrated the opposite pattern: temperature exerted a positive effect and wind did not. Given that cursorial species remain in direct contact with the bare sand, they experience higher temperatures compared to flying species due to the fast and more extreme heating of sandy surfaces. However they do not have specific mechanisms in order to thermoregulate body temperature (Maes et al., 2006; Mutiibwa et al., 2015) and it is thus expected that their movement is more directly linked to the temperatures within their close environment. Our measurements reflect this, on the condition that temperature measured at observer height is directly related to surface temperature. Nevertheless, extremely high temperatures can potentially become detrimental when a certain thermal threshold is reached, and consequently impede any movement (Heinrich, 1995). However, we did not observe such conditions.

The movement direction of both flying and cursorial species exhibited a clear bias towards habitat areas, indicating that the intention of moving towards suitable environments is a more important driver for orientation than wind direction (Acevedo & Fletcher, 2017). This was supported by the data from Kalmthout, where the movement direction should be relatively random due to the proximity of suitable habitats in every direction. At this site, we indeed observed that movement was more random and correlated with wind direction, indicating that the arthropods at De Panne actively chose to move towards suitable habitat. Homing behaviour and site fidelity have been shown in Hymenoptera, Lepidoptera and Blattodea, indicating that insects are indeed able to actively navigate the landscape and determine where suitable areas are located (Boeddeker et al., 2015; Moura et al., 2022; Rivault & Durier, 2004).

Although the direction of movement is clearly related to the location of habitat, movement direction was not completely decoupled from wind conditions. For both flying and cursorial species, strong upwind negatively impacted the returning rates towards the habitat patch, indicating the wind’s ability to divert individuals away from patch boundaries, even if they are able to navigate. Nonetheless, the preference for a specific direction persisted despite wind resistance, implying that arthropods are willing to endure, at least at small spatial scales, the energetic costs associated with navigating towards hospitable patches. Similarly, tailwind assistance could minimize energy expenditure during movement, which is in line with much larger-scale observations from migratory species (Knight et al., 2019; Knoblauch et al., 2021). Only when winds were aligned with the habitat direction, a clear-cut positive correlation between movement and wind direction was observed, indicating that arthropods opportunistically harness winds to facilitate movement towards their intended destinations. Tailwinds are known to drive large-scale migratory events (Åkesson, 2016), but are thus equally important for small-scale movement behaviours at patch boundaries.

Larger flying arthropods moved more towards the habitat compared to smaller individuals and species. A large body size provides advantages in handling upwind movement (Kuenen & Cardé, 1993) and is potentially related to a better-developed perceptual range (i.e. the distance at which an individual can sense its suitable environment). For cursorial arthropods, only phylogenetically uncorrected models showed that smaller taxa are more inclined to move towards suitable habitats. This is most likely related to the abundance of taxa dominated by small species with more extensive perceptual ranges in our study area. For instance, small-bodied arthropod taxa like spiders and ants are known to mainly navigate based on social, chemical and/or tactical cues, which might increase their perceptual range compared to species that solely rely on visual cues (Foelix, 2011).

The tendency of actively reaching a suitable habitat patch (often interpreted as success when costs of entering the matrix are high) depends heavily on both body size and prevailing weather conditions. Larger body sizes and the ability to fly provide advantages for arthropods navigating through the matrix, enhancing movement efficiency. Moreover, ambient temperatures and wind speeds can either augment or impede small-scale movement efficiency, depending on factors like wind direction and thermal thresholds. Thus, movement behaviours at patch boundaries are condition dependent, non-random (Goicolea et al., 2021), and reflect an active decision-making process (Bonte et al., 2012; Clobert et al., 2009; Poethke et al., 2011).

Such non-random movements and dispersal patterns have the strong potential to impact metapopulation dynamics and long-term persistence (Mortier et al., 2019). Therefore, in the context of arthropod metacommunity dynamics, we expect spatial processes and environmental variation to have a greater impact on communities of large-sized flying and smaller, cursorial species (De Bie et al., 2012). Our study provides a mechanistic behavioural perspective on movement at habitat boundaries, which can be integrated in predictive biodiversity models to better understand future metacommunity processes (Urban et al., 2016). This is particularly important given the impending effects of climate change, which are anticipated to alter body sizes (Daly et al., 2024; Davies, 2019; Polidori et al., 2020), temperatures and wind patterns (Greene et al., 2010). This suggests that global warming will not only impact population sizes by increasing stress on a local scale but that it may also directly influence behaviours at habitat boundaries. Therefore, shifting weather patterns could facilitate or constrain movement among habitat patches, directly affecting the composition and future viability of arthropod metacommunities.

## Declarations

### Availability of data and materials

Data available from Zenodo https://doi.org/10.5281/zenodo.8232574 (Logghe et al., 2024).

### Competing interests

The authors declare that they have no competing interests.

### Funding

Garben Logghe was funded by FWO (grantnr: 1130223N). Charlotte Taelman was funded by BOF (grantnr: BOF.24Y.2021.0012.01) and FWO (grantnr: 1SH0I24N).

### Author’s contributions

Garben Logghe, Charlotte Taelman, Florian Van Hecke and Dries Bonte designed the experiments. Charlotte Taelman, Florian Van Hecke and Garben Logghe conducted the practical work. Garben Logghe and Charlotte Taelman analysed the data and wrote the first draft of the manuscript. All authors contributed substantially to the interpretation of the results and revision of the manuscript.

## Acknowledgements

We would like to thank everyone who assisted with the fieldwork: Margaux Pottiez, Ruben Van De Walle, Arthur Van Roey and Elias Van Den Broeck. Special thanks to Johan Lamaire (Agency for Nature and Forests—Flemish Government) and Rudi Delvaux (Grenspark Kalmthoutse Heide) for allowing us access to restricted areas in nature reserves De Westhoek (De Panne) and Kalmthoutse Heide respectively. Finally, we would like to thank three anonymous reviewers for their valuable comments on a previous version of the manuscript.

## Notes

https://doi.org/10.5281/zenodo.8232574

### Competing Interest Statement

The authors have declared no competing interest.

### Summary of Updates

Added an ethics statement to the methodology; revised the position of Figure 1

