## Supplementary material for "Unravelling arthropod movement in natural landscapes: small-scale effects of body size and weather conditions"

#### S.1 Sampling locations for the experimental fieldwork

A) De Panne site

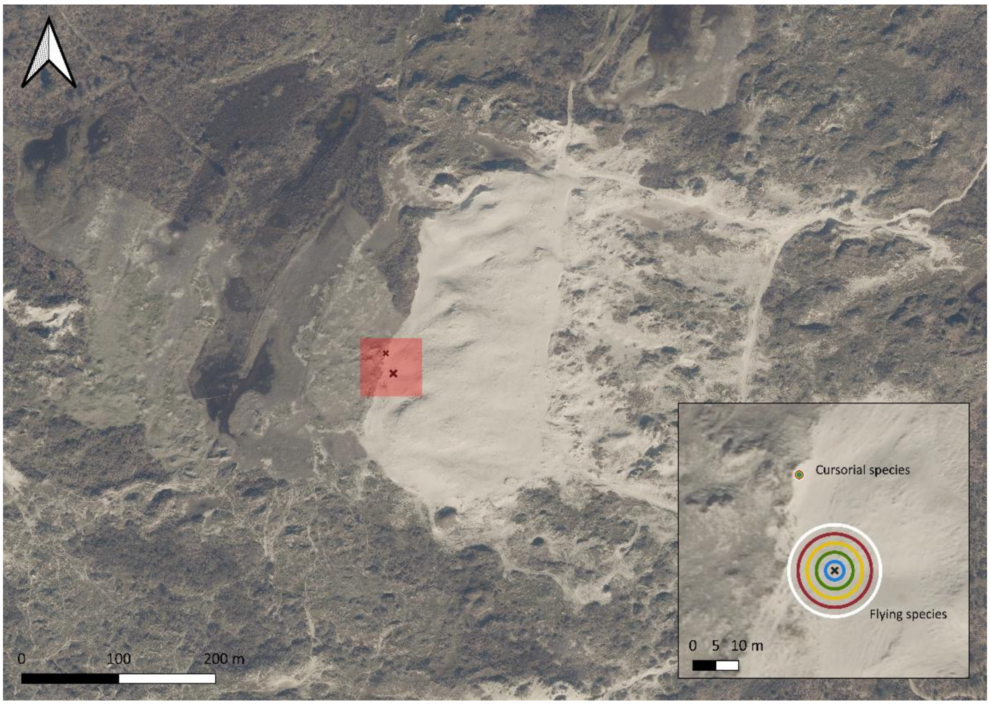


B) Kalmthout site

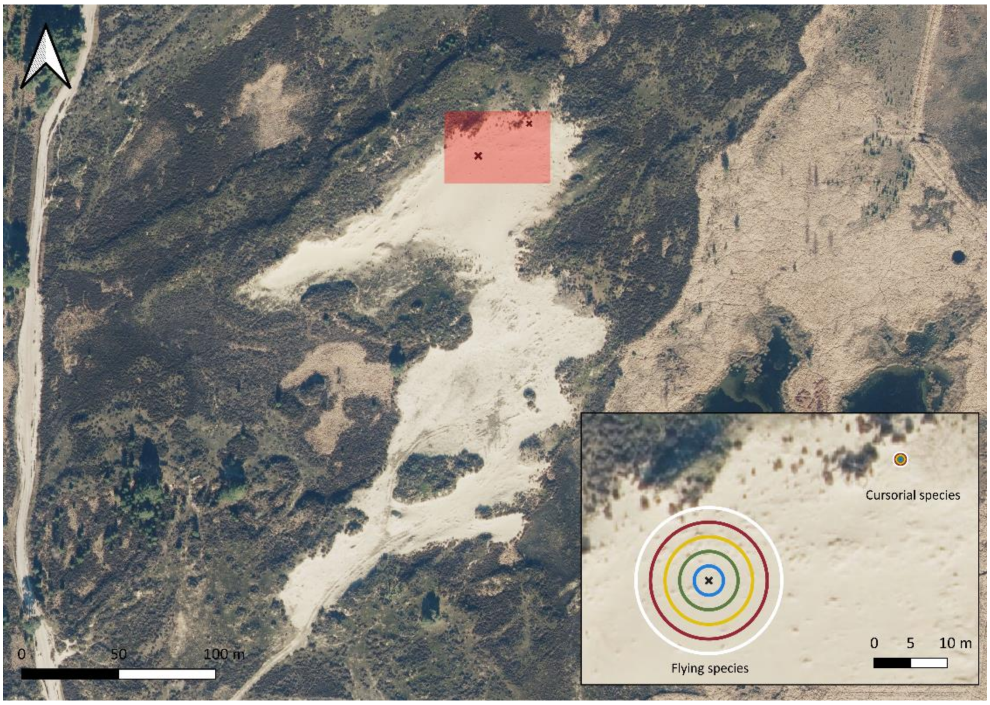


*Figure S.1: The two sampling sites in Belgium. A) Open sand of grey dunes bordering dune grassland and shrubland. B) Open sand surrounded by dry and wet heathland patches. Black crosses indicate the location of the experimental arenas.*

#### S.2 Overview of sampled species

*Table S.2: Overview of the number of families, species and individuals that were tested for each arthropod order.*

| ORDER | FAMILIES | SPECIES | INDIVIDUALS |
|---|---|---|---|
| **ARANEAE** | 15 | 46 | 85 |
| **BLATTODEA** | 1 | 1 | 9 |
| **COLEOPTERA** | 12 | 28 | 102 |
| **DERMAPTERA** | 1 | 1 | 1 |
| **DIPTERA** | 13 | 27 | 234 |
| **HEMIPTERA** | 10 | 53 | 144 |
| **HYMENOPTERA** | 16 | 49 | 285 |
| **ISOPODA** | 2 | 3 | 40 |
| **LEPIDOPTERA** | 11 | 30 | 161 |
| **NEUROPTERA** | 1 | 1 | 1 |
| **ODONATA** | 3 | 8 | 35 |
| **OPILIONES** | 1 | 4 | 47 |
| **ORTHOPTERA** | 3 | 14 | 56 |
| **TRICHOPTERA** | 1 | 1 | 1 |

### S.3 Principal Component Analysis

*Table S.3: Output of the PCA for both flying and cursorial individuals.*

| | PC1 | PC2 | PC3 | PC4 |
|---|---|---|---|---|
| **Flying locomotion** | | | | |
| Standard deviation | 1.21 | 1.02 | 0.94 | 0.78 |
| Body size | 0.10 | -0.87 | -0.45 | 0.18 |
| Temperature | -0.62 | -0.15 | 0.42 | 0.64 |
| Average wind speed | 0.65 | 0.24 | -0.03 | 0.72 |
| Movement speed | 0.42 | -0.41 | 0.79 | -0.21 |
| **Cursorial locomotion** | | | | |
| Standard deviation | 1.17 | 1.06 | 0.93 | 0.81 |
| Body size | -0.08 | -0.78 | 0.51 | 0.35 |
| Temperature | 0.67 | -0.17 | 0.29 | -0.66 |
| Average wind speed | -0.60 | -0.41 | -0.29 | -0.63 |
| Movement speed | 0.44 | -0.44 | -0.75 | 0.22 |

### S.4 Correlation between movement speed and other factors

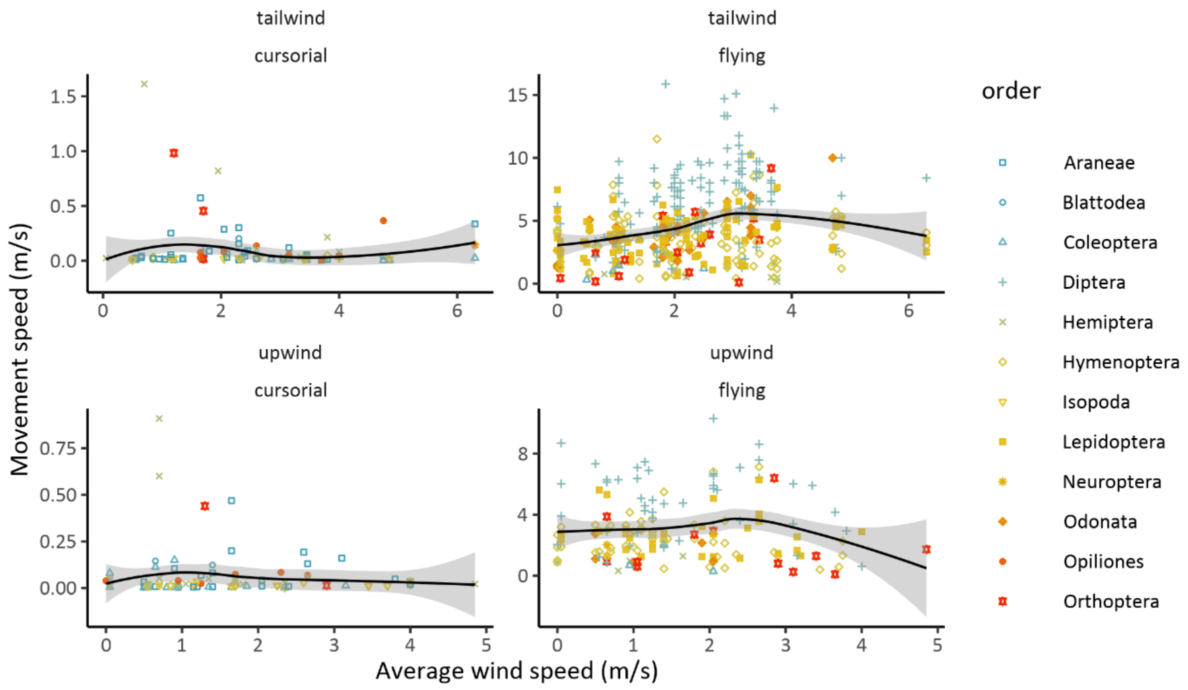


*Figure S.4.1: The effect of the wind speed on the movement speed of cursorial and flying individuals, separated in upwind and tailwind conditions.*

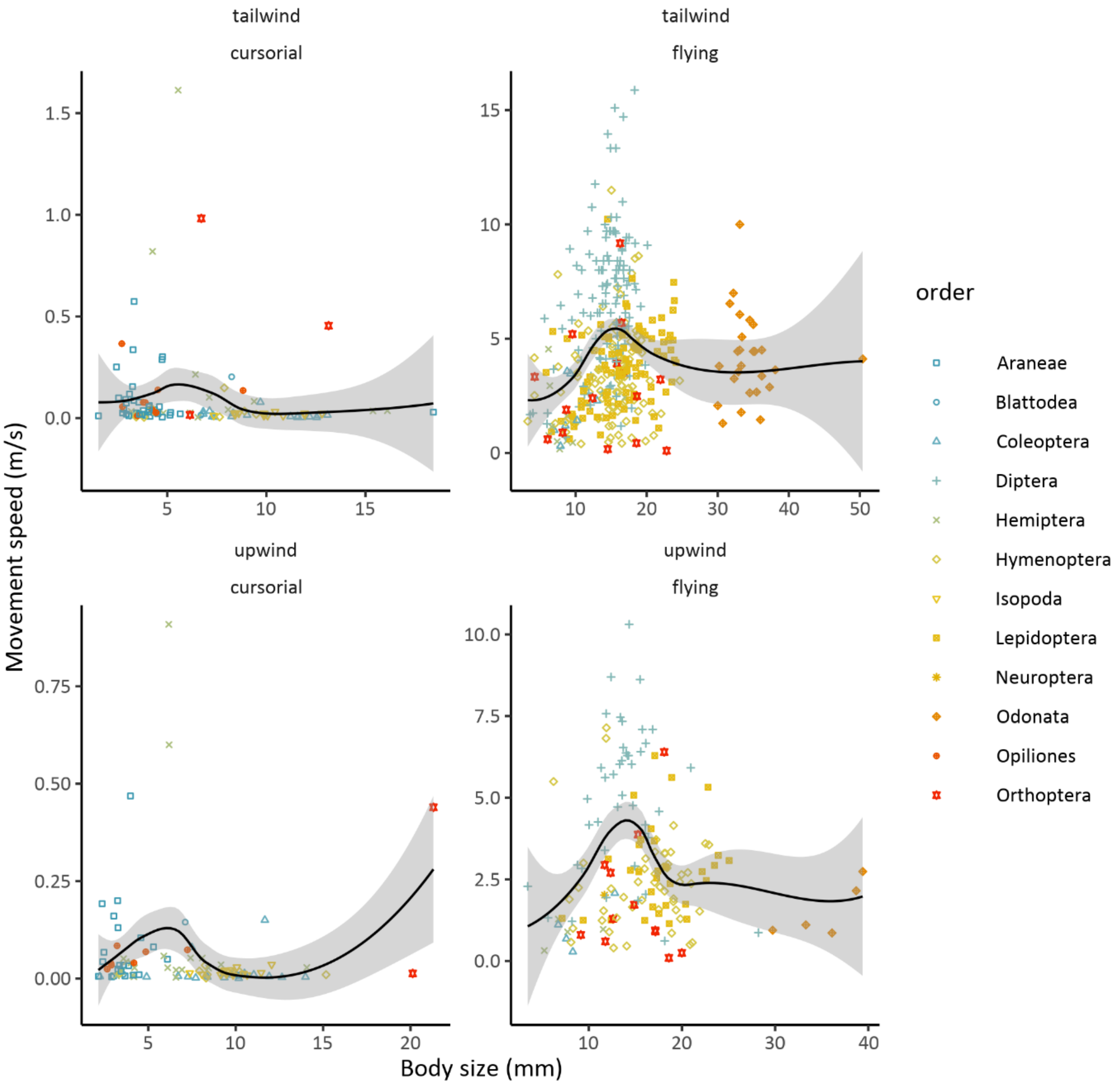


*Figure S.4.2: The effect of the body size on the movement speed of cursorial and flying individuals, separated in upwind and tailwind conditions.*

### S.5 Bayesian Generalized Linear Models

*Table S.5.1: Results from the Bayesian models with movement speed as the dependent variable and Gaussian distribution. Effect sizes were deemed clearly determined when 0 was not included in the 95% credibility intervals of the posteriors (indicated by *).*

| | Posterior mean | 95% confidence interval |
|---|---|---|
| **Flying speed (all data)** | | |
| Intercept | 3.85 | [3.65; 4.04] * |
| Body size | 0.25 | [0.06; 0.44] * |
| Temperature | -0.09 | [-0.29; 0.65] |
| Wind speed | 0.45 | [0.25; 0.65] * |
| ***Phylogenetic correction*** | | |
| Intercept | 2.77 | [-0.29; 5.67] |
| Body size | 0.36 | [0.10; 0.61] * |
| Temperature | -0.04 | [-0.20; 0.12] |
| Wind speed | 0.36 | [0.20; 0.52] * |
| **Flying speed (tailwind)** | | |
| Intercept | 4.41 | [4.12; 4.70] * |
| Body size | 0.25 | [0.00; 0.51] |
| Temperature | -0.07 | [-0.40; 0.27] |
| Wind speed | 0.54 | [0.24; 0.83] * |
| ***Phylogenetic correction*** | | |
| Intercept | 3.28 | [0.19; 6.23] * |
| Body size | 0.57 | [0.20; 0.93] * |
| Temperature | -0.19 | [-0.44; 0.07] |
| Wind speed | 0.37 | [0.13; 0.60] * |
| **Flying speed (upwind)** | | |
| Intercept | 3.05 | [2.69; 3.42] * |
| Body size | -0.18 | [-0.54; 0.18] |
| Temperature | -0.21 | [-0.58; 0.15] |
| Wind speed | -0.16 | [-0.59; 0.28] |
| ***Phylogenetic correction*** | | |
| Intercept | 2.01 | [-0.88; 4.99] |
| Body size | 0.17 | [-0.26; 0.59] |
| Temperature | -0.04 | [-0.20; 0.12] |
| Wind speed | 0.01 | [-0.28; 0.30] |
| **Cursorial speed (all data)** | | |
| Intercept | 0.09 | [0.07; 0.10] * |
| Body size | 0.01 | [-0.01; 0.02] |
| Temperature | 0.04 | [0.02; 0.05] * |
| Wind speed | -0.00 | [-0.02; 0.01] |
| ***Phylogenetic correction*** | | |
| Intercept | 0.12 | [-0.19; 0.44] |
| Body size | -0.00 | [-0.02; 0.02] |
| Temperature | 0.03 | [0.01; 0.04] * |
| Wind speed | -0.01 | [-0.02; 0.01] |
| **Cursorial speed (tailwind)** | | |
| Intercept | 0.10 | [0.05; 0.14] * |
| Body size | -0.02 | [-0.06; 0.02] |
| Temperature | 0.03 | [-0.01; 0.07] |
| Wind speed | -0.00 | [-0.04; 0.03] |

| | | |
|---|---|---|
| ***Phylogenetic correction*** | | |
| Intercept | 0.12 | [-0.14; 0.40] |
| Body size | -0.03 | [-0.08; 0.02] |
| Temperature | 0.02 | [-0.02; 0.06] |
| Wind speed | -0.01 | [-0.04; 0.03] |
| **Cursorial speed (upwind)** | | |
| Intercept | 0.05 | [0.02; 0.08] * |
| Body size | -0.00 | [-0.03; 0.02] |
| Temperature | 0.02 | [-0.00; 0.05] |
| Wind speed | -0.00 | [-0.03; 0.03] |
| ***Phylogenetic correction*** | | |
| Intercept | 0.06 | [-0.19; 0.35] |
| Body size | 0.01 | [-0.02; 0.04] |
| Temperature | 0.02 | [-0.00; 0.05] |
| Wind speed | 0.00 | [-0.02; 0.03] |

*Table S.5.2: Results from the Bayesian models with movement direction as the dependent variable and binomial distribution with logit link function. Effect sizes were deemed clearly determined when 0 was not included in the 95% credibility intervals of the posteriors (indicated by *).*

| | Posterior mean | 95% confidence interval |
|---|---|---|
| **Flying navigation (all data)** | | |
| Intercept | 0.22 | [0.05; 0.39] * |
| Body size | 0.26 | [0.09; 0.44] * |
| Wind speed | -0.26 | [-0.44; -0.09] * |
| ***Phylogenetic correction*** | | |
| Intercept | 0.09 | [-0.75; 0.64] |
| Body size | 0.27 | [0.08; 0.45] * |
| Wind speed | -0.27 | [-0.45; -0.09] * |
| **Flying navigation (tailwind)** | | |
| Intercept | 0.25 | [0.01; 0.50] * |
| Body size | 0.14 | [-0.09; 0.40] |
| Wind speed | -0.19 | [-0.42; 0.04] |
| ***Phylogenetic correction*** | | |
| Intercept | -0.12 | [-1.88; 1.49] |
| Body size | 0.16 | [-0.11; 0.44] |
| Wind speed | -0.22 | [-0.47; 0.03] |
| **Flying navigation (upwind)** | | |
| Intercept | 0.11 | [-0.32; 0.54] |
| Body size | 0.30 | [-0.11; 0.72] |
| Wind speed | -0.94 | [-1.45; -0.47] * |
| ***Phylogenetic correction*** | | |
| Intercept | -0.19 | [-2.33; 1.69] |
| Body size | 0.26 | [-0.22; 0.77] |
| Wind speed | -1.08 | [-1.66; -0.54] * |
| **Cursorial navigation (all data)** | | |
| Intercept | 0.35 | [0.12; 0.58] * |
| Body size | -0.26 | [-0.50; -0.02] * |
| Wind speed | -0.09 | [-0.32; 0.14] |
| ***Phylogenetic correction*** | | |
| Intercept | 0.31 | [-0.97; 1.52] |

| | | |
|---|---|---|
| Body size | -0.07 | [-0.37; 0.25] |
| Wind speed | -0.14 | [-0.39; 0.10] |
| **Cursorial navigation (tailwind)** | | |
| Intercept | 0.59 | [0.18; 1.01] * |
| Body size | -0.22 | [-0.64; 0.22] * |
| Wind speed | -0.02 | [-0.41; 0.36] |
| ***Phylogenetic correction*** | | |
| Intercept | 0.58 | [-0.31; 1.45] |
| Body size | -0.18 | [-0.70; 0.34] |
| Wind speed | -0.02 | [-0.39; 0.36] |
| **Cursorial navigation (upwind)** | | |
| Intercept | 0.21 | [-0.25; 0.67] |
| Body size | -0.48 | [-0.98; -0.02] * |
| Wind speed | -0.68 | [-1.18; -0.20] * |
| ***Phylogenetic correction*** | | |
| Intercept | 0.18 | [-1.11; 1.44] |
| Body size | -0.60 | [-1.27; -0.01] * |
| Wind speed | -0.69 | [-1.24; -0.18] * |

### S.6 Rayleigh tests

*Table S.6: Output for the Rayleigh's test of uniformity for different orders, wind directions and locomotion modes in both sampling sites. P-values are corrected for multiplicity using the Benjamini-Hochberg method. The significance level is indicated with ***. Theta (ϑp) represents the mean movement direction (in radians), together with the circular standard deviation on the average movement (SD, also in radians). R is the mean resultant length and is a measure of the degree of bias in movement. When R is 0, movement is uniformly distributed on a circle. When R is equal to 1, all movement is biased in one particular direction. Datasets that consisted of less than 8 observations, were not considered in this analysis, as the bias in a certain direction (8 possible directions) would otherwise not be correctly notified.*

| Order | θ | sd | R | p-value | | n | Location |
|---|---|---|---|---|---|---|---|
| Lepidoptera | -16.03 | 1.38 | 0.39 | 0.00 | *** | 52 | Kalmthout |
| Diptera | 109.47 | 1.71 | 0.23 | 0.51 | | 13 | Kalmthout |
| Hymenoptera | -93.34 | 1.83 | 0.19 | 0.19 | | 47 | Kalmthout |
| Odonata | -42.26 | 1.05 | 0.57 | 0.00 | *** | 32 | Kalmthout |
| Isopoda | -59.35 | 1.73 | 0.23 | 0.15 | | 38 | De Panne |
| Opiliones | 0.00 | 1.44 | 0.36 | 0.03 | *** | 27 | De Panne |
| Orthoptera | 16.32 | 1.72 | 0.23 | 0.09 | | 47 | De Panne |
| Coleoptera | -134.61 | 2.15 | 0.10 | 0.49 | | 75 | De Panne |
| Hemiptera | -11.87 | 1.24 | 0.46 | 0.00 | *** | 82 | De Panne |
| Lepidoptera | -33.95 | 1.41 | 0.37 | 0.00 | *** | 103 | De Panne |
| Araneae | -1.67 | 1.29 | 0.43 | 0.00 | *** | 89 | De Panne |
| Diptera | -29.21 | 1.47 | 0.34 | 0.00 | *** | 202 | De Panne |
| Hymenoptera | -28.59 | 1.71 | 0.23 | 0.00 | *** | 200 | De Panne |

| Locomotion | θ | sd | R | p-value | | n | Location |
|---|---|---|---|---|---|---|---|
| flying | -37.32 | 1.62 | 0.27 | 0.00 | *** | 146 | Kalmthout |
| flying | -26.50 | 1.54 | 0.30 | 0.00 | *** | 560 | De Panne |
| cursorial | -12.99 | 1.66 | 0.25 | 0.00 | *** | 313 | De Panne |

| Wind | θ | sd | R | p-value | | n | Location |
|---|---|---|---|---|---|---|---|
| E | -44.70 | 1.80 | 0.20 | 0.15 | | 49 | Kalmthout |
| S | 96.18 | 1.81 | 0.19 | 0.60 | | 14 | Kalmthout |
| NE | -45.85 | 1.28 | 0.44 | 0.00 | *** | 75 | Kalmthout |
| SE | 64.47 | 1.13 | 0.53 | 0.10 | | 8 | Kalmthout |
| N | -68.44 | 1.50 | 0.32 | 0.00 | *** | 312 | De Panne |
| E | -5.30 | 1.38 | 0.38 | 0.00 | *** | 173 | De Panne |
| S | 37.74 | 1.43 | 0.36 | 0.00 | *** | 79 | De Panne |
| W | 12.94 | 1.26 | 0.45 | 0.00 | *** | 38 | De Panne |
| NE | -31.87 | 1.32 | 0.42 | 0.00 | *** | 88 | De Panne |
| SE | 23.67 | 0.83 | 0.71 | 0.00 | *** | 11 | De Panne |
| NW | -1.16 | 1.39 | 0.38 | 0.00 | *** | 92 | De Panne |
| SW | 5.24 | 1.83 | 0.19 | 0.05 | | 82 | De Panne |

| Location (all data) | θ | sd | R | p-value | | n |
|---|---|---|---|---|---|---|
| Kalmthout | -37.32 | 1.62 | 0.27 | 0.00 | *** | 146 |
| De Panne | -22.22 | 1.58 | 0.29 | 0.00 | *** | 875 |

### S.7 Jammalamadaka-Sarma correlations

*Table S.7: Output for the Jammalamadaka-Sarma correlation tests for the different taxonomic orders and locomotion modes in both sampling locations (n = number of samples). P-values are corrected for multiplicity using the Benjamini-Hochberg method. The significance level is indicated with ***. The correlation coefficient is a circular version of Pearson's product moment correlation. The test statistic represents the value of the statistic to test the null hypothesis that the correlation coefficient is 0. The p-value is calculated using a large-sample normal approximation. If two circular variables are completely independent from each other, then the correlation coefficient is 0. However, the opposite does not always hold; when the correlation coefficient is 0 this does not imply that two circular random variables are independent. Datasets that consisted of less than 8 observations, were not considered in this analysis, as not all possible movement and wind directions would be represented.*

| Order | Correlation | Test statistic | p-value | | n | Location |
|---|---|---|---|---|---|---|
| Lepidoptera | 0.36 | 2.50 | 0.01 | *** | 52 | Kalmthout |
| Diptera | -0.06 | -0.23 | 0.82 | | 13 | Kalmthout |
| Hymenoptera | -0.23 | -1.49 | 0.14 | | 47 | Kalmthout |
| Odonata | 0.31 | 1.52 | 0.13 | | 32 | Kalmthout |
| Isopoda | -0.27 | -1.68 | 0.09 | | 38 | De Panne |
| Opiliones | 0.28 | 1.69 | 0.09 | | 27 | De Panne |
| Orthoptera | -0.19 | -1.30 | 0.19 | | 47 | De Panne |
| Coleoptera | -0.14 | -1.20 | 0.23 | | 75 | De Panne |
| Hemiptera | -0.22 | -2.08 | 0.04 | | 82 | De Panne |
| Lepidoptera | -0.23 | -2.42 | 0.02 | | 103 | De Panne |
| Araneae | 0.34 | 3.29 | 0.00 | *** | 89 | De Panne |
| Diptera | 0.19 | 2.61 | 0.01 | *** | 202 | De Panne |
| Hymenoptera | 0.10 | 1.35 | 0.18 | | 200 | De Panne |

| Locomotion | Correlation | Statistic | p-value | n | Location |
|---|---|---|---|---|---|
| flying | 0.14 | 1.64 | 0.10 | 146 | Kalmthout |
| flying | 0.09 | 2.05 | 0.04 | 560 | De Panne |
| cursorial | 0.10 | 1.85 | 0.06 | 313 | De Panne |

| Location (all data) | Correlation | Test statistic | p-value | n |
|---|---|---|---|---|
| Kalmthout | 0.14 | 1.64 | 0.10 | 146 |
| De Panne | 0.07 | 2.08 | 0.04 | 875 |

### S.8 Circular bar plots of movement direction

#### A) De Panne site

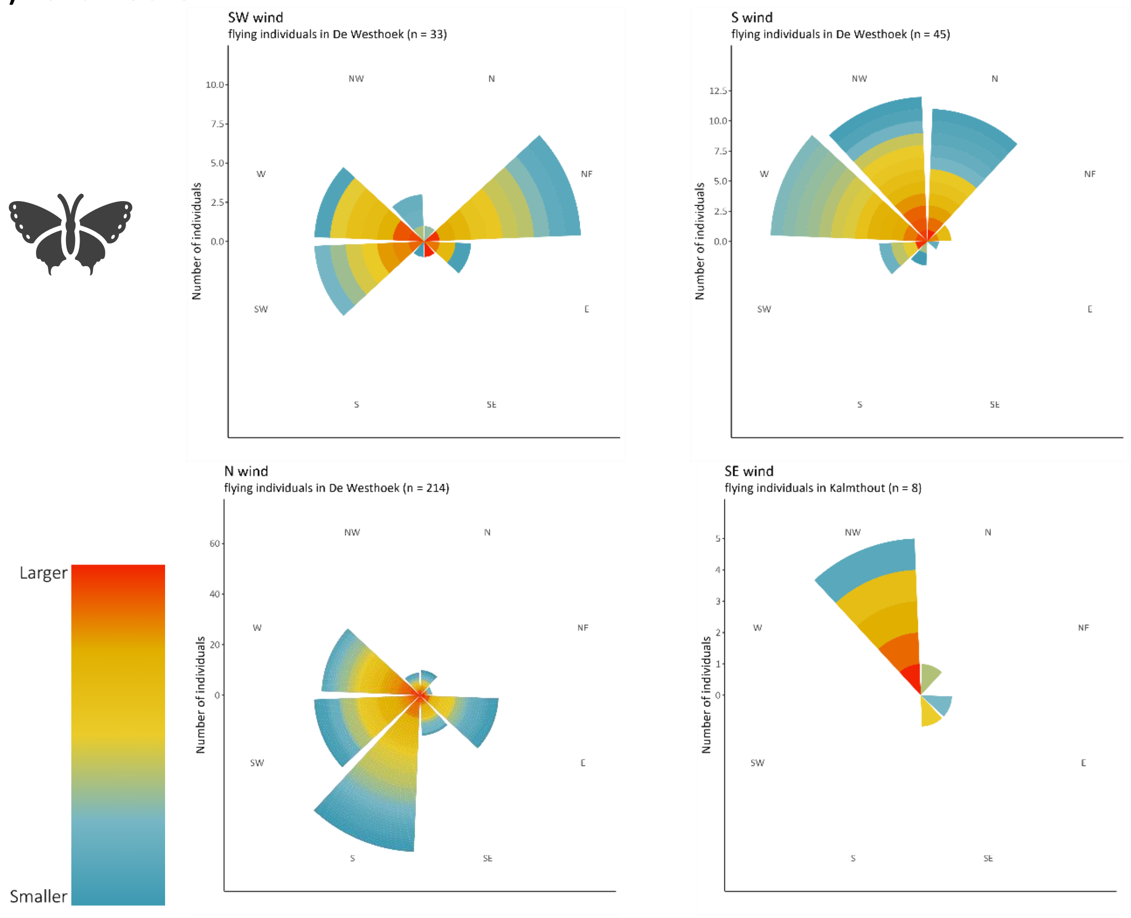

#### B) Kalmthout site

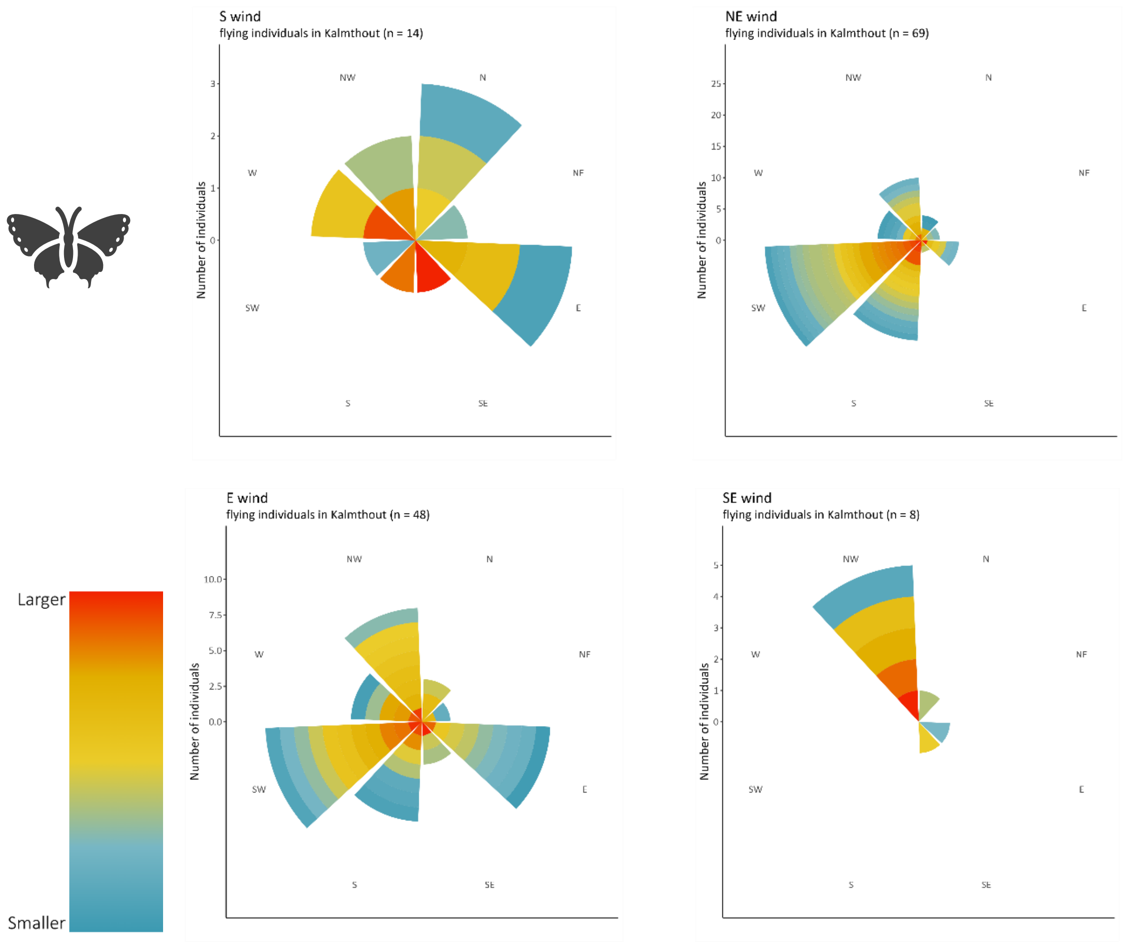


*Fig. S.8.1: Circular bar plots of movement directions for flying species under the different wind direction scenarios in both sampling sites. Each slice on the bars represents an individual, which is colored according to its body size.*

#### A) De Panne site

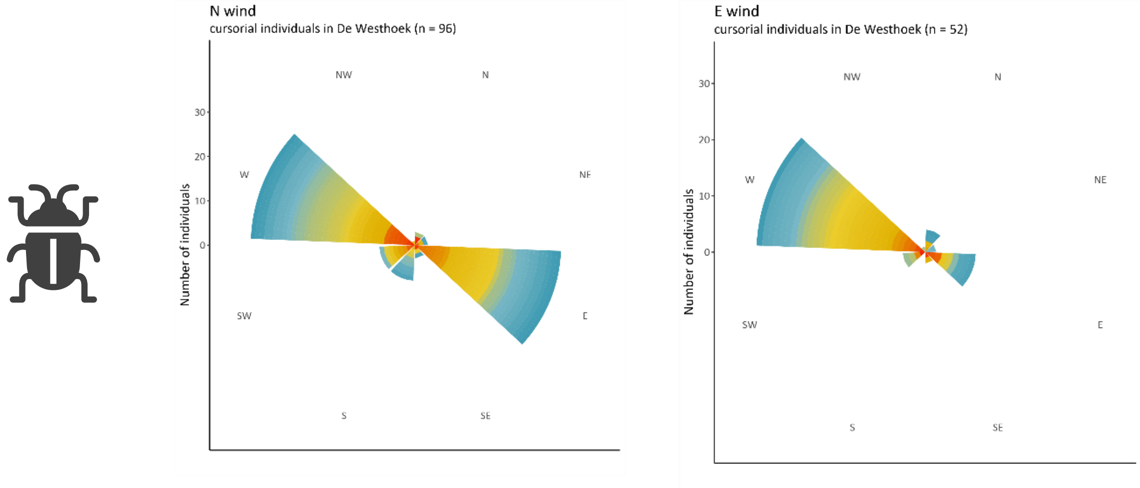

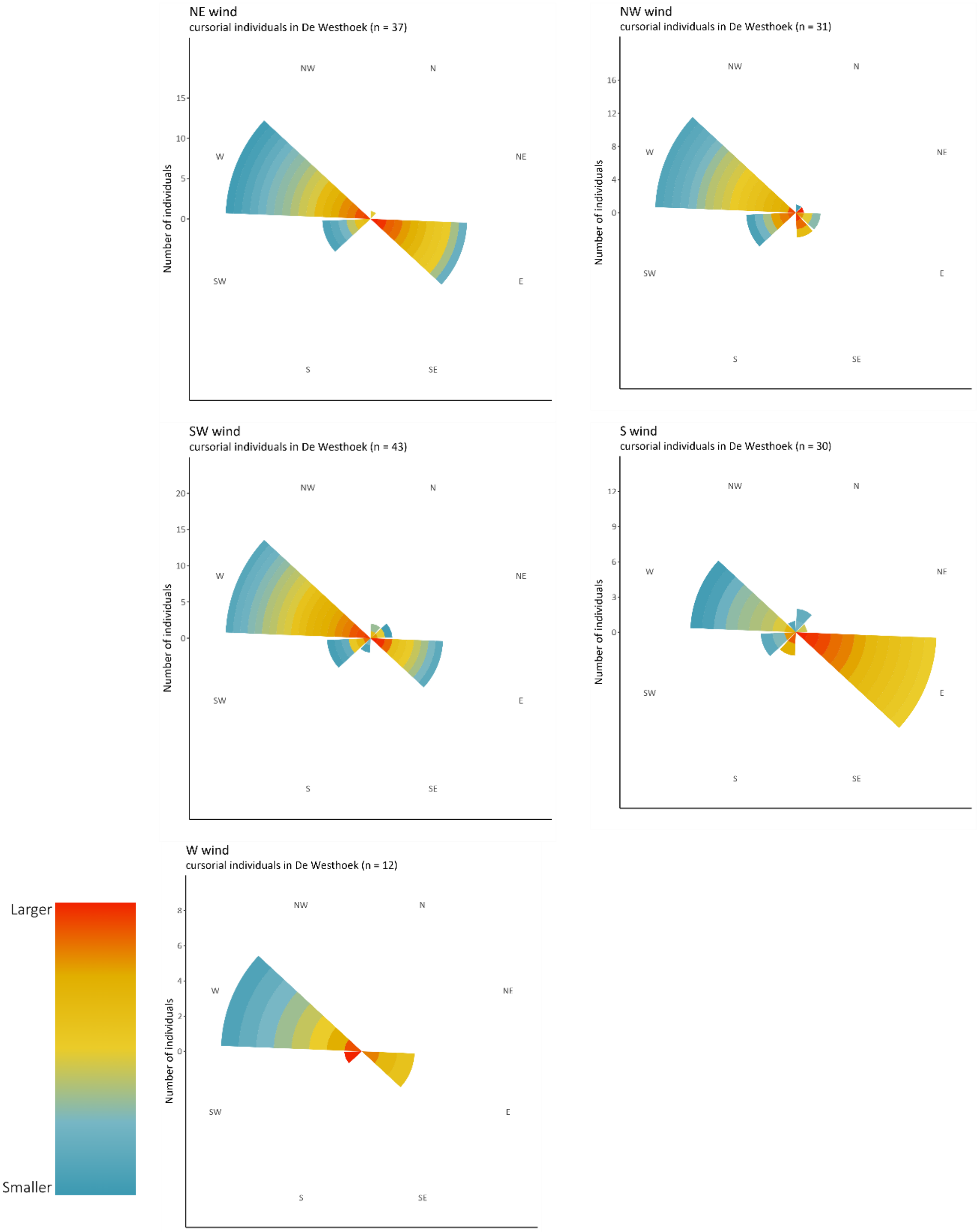


*Fig. S.8.2: Circular bar plots of movement directions for cursorial species under 7 different wind direction scenarios in De De Panne (Kalmthout site had too few observations and are thus left out). Each slice on the bars represents an individual, which is colored according to its body size.*

### S.9 Wind speed data per site

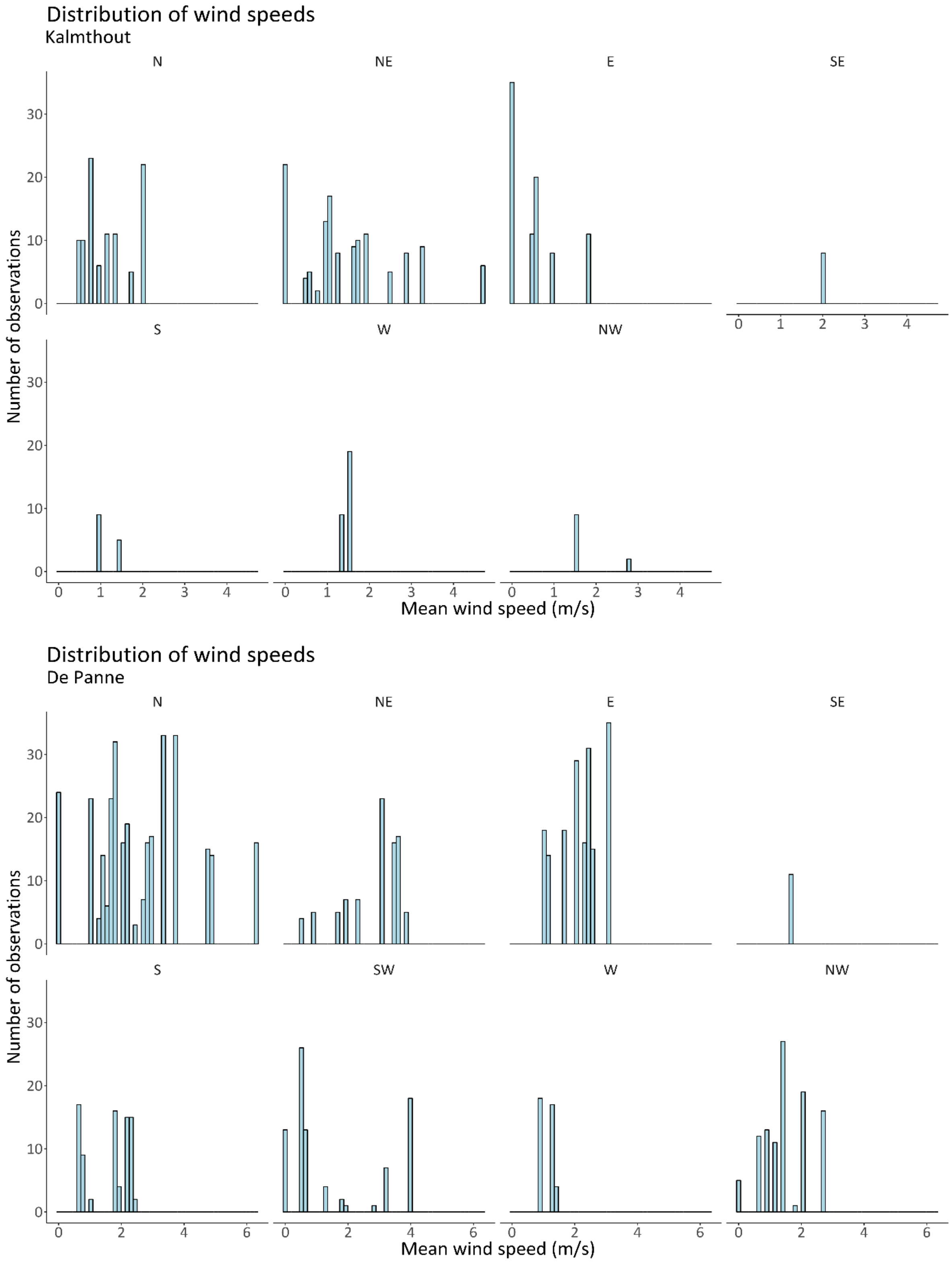


*Fig. S.9: Histograms showing the frequency of certain wind speeds at the two field sites. The height of the bars indicate the number of times a certain wind speed was measured for each wind direction.*
